# Insertion of rare autism variants in synaptic genes induce novel behavioral phenotypes in *C. elegans*

**DOI:** 10.64898/2025.12.19.695228

**Authors:** Dustin Haskell, William R. Haury, Myra Granato, Brandon L. Bastien, Michael P. Hart

## Abstract

Neurodevelopmental conditions and disorders, including autism, involve a complex interplay of genetic, environmental, and developmental factors. Despite this complexity, genetic studies have identified more than 150 candidate genes that increase risk for autism and related neurodevelopmental and neuropsychiatric conditions. Unsurprisingly, synaptic genes are a large proportion of these genes, likely due to their roles in the formation and maintenance of synaptic architecture, function, and the plasticity of neurons and circuits. The association of synaptic genes with autism and similar conditions is driven by all types of genetic variation, including inherited and *de novo* rare variants that have unknown impacts on the function of the gene. Here we insert 4 conserved rare variants in the *C. elegans* orthologs of *NLGN4X, NRXN1*, and *SHANK3,* and define their impact on gene function compared to known loss of function variants using behavioral assays. We find that the rare variants impact multiple foraging behaviors, with each gene and variant having a unique pattern of behavioral changes and functional impact. The NLGN4X(A283T) variant induced clear loss of function, while NLGN4X(G84R) induces a loss of function in one behavior, but a gain of function in another behavior. The NRXN1(L18Q) variant induced remarkable loss and gain of functions with distinct impacts across each behavior. The SHANK3(L143P) variant induced partial loss of function in a single behavior. We also identify for the first time that loss of function of *shn-1/SHANK3* alters social feeding and food response behaviors. We uncover a remarkably complex impact of rare variants in synaptic genes, with differential impacts across behaviors, highlighting the importance of broad behavioral analysis and the nuanced effects of missense variants compared to loss of function alleles. Together, we define the complex functional impact of each variant on gene function, compare the impact of variants and genes across multiple behaviors, and provide further support for the use of *C. elegans* to define the impact of genetic variation derived from human neurodevelopmental and neuropsychiatric disorders.

## INTRODUCTION

Many neurodevelopmental and neuropsychiatric conditions and disorders are defined and diagnosed by characteristic changes in behavior. For example, autism is clinically characterized by changes in social and communication behaviors and the presence of repetitive or restrictive behaviors^1^. However, for many on the autism spectrum, additional changes in genetics^2–4^, neuropsychiatric and psychiatric features^5,6,7,8^ can be involved, and behavioral changes can range in magnitude or severity amongst individuals^9,10^ and show sex variation^11^. Therefore, the vast variation observed in autism, related conditions, and neurodiversity, has limited our understanding of the developmental and molecular mechanisms that underlie the spectrum of behavioral changes.

Recent case-control, familial, and large cohort genetic and genomic studies, have identified hundreds of genes associated with autism, including many synaptic genes in the list of highest confidence risk genes^12,13^. Within autism-associated genes, perturbations can include deletions (both large and small indels), missense, nonsense, and other types of variants. While many mutation types, such as large deletions, have a well-defined impact on the gene by the nature of the variant itself, others often have unknown impact or significance. When regarding these rare variants, particularly missense variants, a number of questions should be considered: 1) do variants impact the gene function, and if so, how do they alter the protein structure and function, 2) do alterations in protein function result in neuronal differences leading to behavioral changes, 3) do variants in different genes impact the same behaviors and can variants in the same gene differentially impact multiple behaviors?

An clear example of this genetic complexity are neurexin genes (*NRXN1-3*)^14,15^ and their canonical binding partners neuroligins (*NLGN1-4*(XY))^16–18^. Variants in neurexins are repeatedly and strongly associated with autism and other syndromes and conditions, including Pitt-Hopkins-like syndrome 2, Tourette syndrome, developmental delay, intellectual disability, and epilepsy (SFARI GENE)^19–22^. The association of *NRXN1* with autism and related conditions is largely driven by large deletions of exons, there are also intronic deletions, missense, and nonsense variants observed with unknown impact. Neurexins play important roles in synaptic organization^23–26^, influencing neuronal dynamics and circuit connectivity^27–29^, and more recently in neuronal plasticity responses^30–32^. Furthermore, neurexins influence the structure and properties of individual synaptic connections, helping maintain excitatory/inhibitory balance within and across circuits. The canonical binding partners of neurexins, neuroligins, are also genetically associated with autism and related conditions and syndromes^18^. The association of neuroligins are primarily driven by missense variants, and the neuroligin proteins are implicated in establishing pre- and post-synaptic polarity through cooperative organization of synaptic architecture, and clustering receptors and other organizing molecules, including neurexins.

Also associated in genetic studies of autism, are shank genes (*SHANK1-3*), which act as critical intracellular scaffolding proteins in post-synaptic densities. *SHANK3* is associated with Phelan-McDermid syndrome, Rett syndrome-like phenotype, Pediatric Acute-Onset Neuropsychiatric Syndrome, developmental delay and intellectual disability (SFARI GENE)^33,33–37^. Broadly, these scaffolding proteins utilize multiple and diverse domains to drive interactions between many post-synaptic molecules. Ankyrin repeat domains interact with SHARPIN and thus indirectly scaffold with cytoskeletal elements^38^. PDZ domains have been shown to directly interact with several subunits of AMPA receptors during the formation of dendritic spines^39^, and SAM domains self-multimerize and direct localization to post-synaptic densities^40^. Taken together, these studies suggest that by anchoring synaptic proteins, especially receptors, to the intracellular surface of synaptic membranes Shank proteins are able to drive the organization of post-synaptic densities. Together, the significant associations of these synaptic genes suggest an important role for synaptic and circuit organization for autism, related disorders, and the behavioral changes associated with each.

To better understand neurodevelopment and the mechanisms and implications of synaptic gene dysfunction on behaviors, recent efforts have aimed to explore the role of genes and variants on neurons, circuits, and behaviors^41–43^. *C. elegans* has emerged as a tractable tool, as they display simple, robust, and stereotyped behaviors that are the result of intact circuits, and where the impact of a gene can be mechanistically defined at the level of single gene isoforms acting in single neurons. In recent work, we described roles for multiple conserved autism-associated genes in *C. elegans* circuits and behaviors. Specifically, two well characterized foraging behaviors in *C. elegans*, the response to food (and food deprivation) and social feeding behaviors^30,44,45^. We found that the alpha and gamma isoforms of the singular neurexin gene in *C. elegans*, *nrx-1,* are required for a hyperactive behavioral response to food deprivation^45^. Further, we found that the long isoform of *nrx-1* is required for a social feeding behavior, but not for a solitary feeding behavior. We found that loss of *nlg-1* had no impact on the response of animals to food or loss of food, it reduced social feeding behavior, with no impact on solitary feeding behavior^30^. In addition to the roles of these genes and the loss of their function, we were able to define the impact of this loss on specific neurons, synaptic connections, and signaling pathways^30,45^

Here we examine the functional and behavioral impact of 4 rare variants from autism probands in *NRXN1*, *NLGN4X*, and *SHANK3* by identifying conserved residues in the respective *C. elegans* gene, *nrx-1*, *nlg-1*, and *shn-1* (**Figure 1A**). We inserted each conserved variant into the respective *C. elegans* orthologous gene using CRISPR Cas9 genome editing. We then tested the impact of each variant on multiple behaviors, including the behavioral response to food and food deprivation, and solitary and social feeding behaviors. Consistent with previous analysis of null or strong loss of function alleles, we observed a significant reduction in activity upon food deprivation in select variants in *nlg-1* and *nrx-1*. This reduction is also present in several variants of *nlg-1* and *nrx-1* even in the presence of food, indicating novel variant-specific phenotypes, and potential gain of function impacts. When examining social feeding behavior, we observed that specific *nrx-1* and *nlg-1* variants decreased aggregate social feeding, similar to loss of function deletion alleles in these genes. We identify a novel role for the *shn-1* in social feeding behavior, where both the loss of function deletion the rare variant decrease social feeding behavior, with the rare variant having a larger impact. Remarkably, while we observe no impact for most genes and variants on solitary feeding behavior, the *nrx-1(sy869)* variant significantly induces aggregate feeding behavior in a solitary background. Together, we identify complex functional impacts of rare variants across multiple synaptic genes, including loss and gain of functions that are shared and differ across each gene and behavior. This work highlights the utility of *C. elegans* to define the behavioral and functional impact of variants in multiple genes by comparing controls and loss of function alleles, and provides additional proof of concept for using *C. elegan*s to model and define rare variants found in complex human conditions.

**Figure 1.**
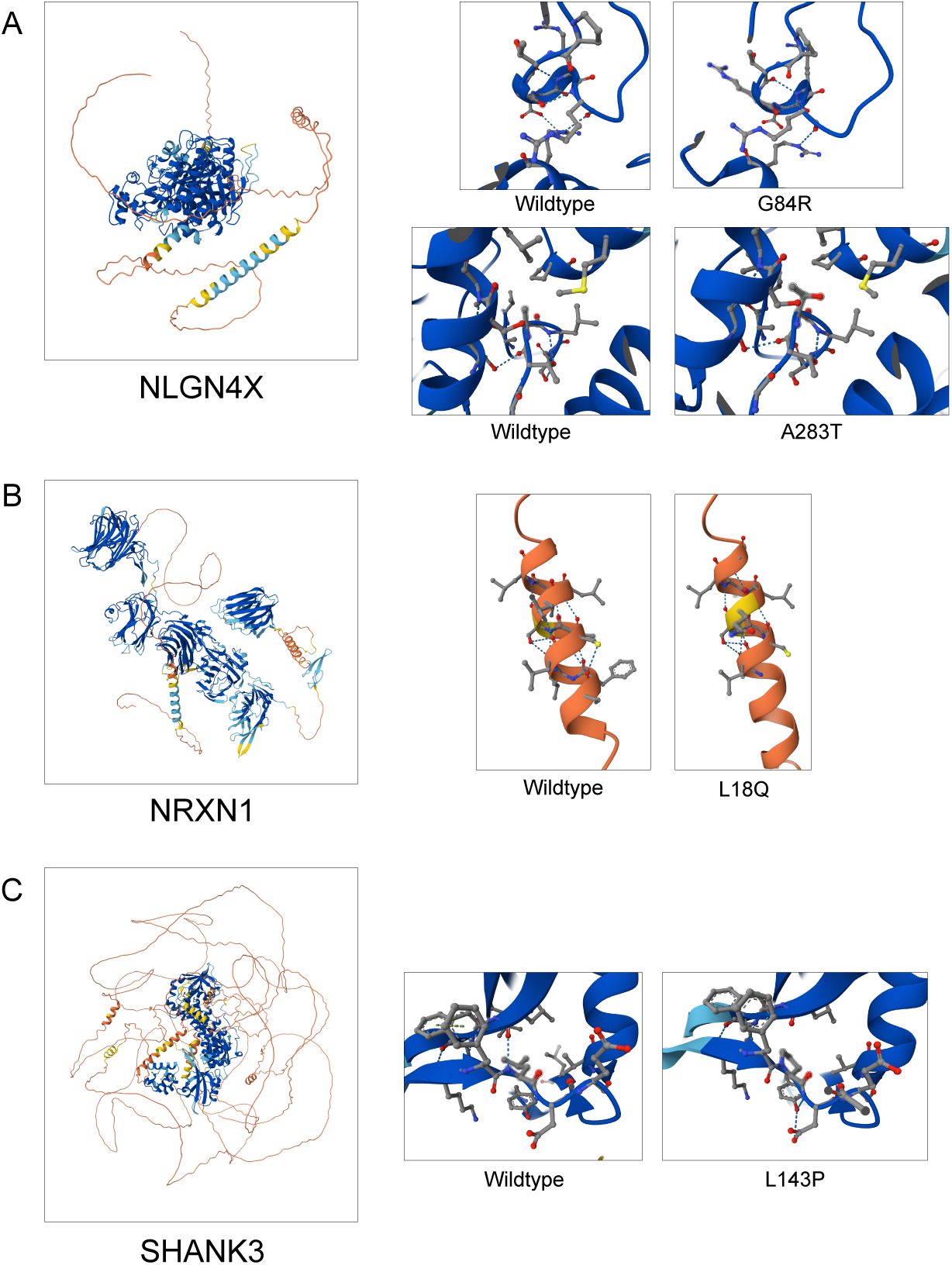
Predicted structural models of synaptic proteins and rare variants. **(A)** NGLN4X structural prediction and predictions of the structural effects of G84R and A283T variations. **(B)** NRXN1 structural prediction and predictions of the structural effects of the L18Q variation. **(C)** SHANK3 structural prediction and predictions of the structural effects of the L143P variation.

## METHODS

### Strain Construction and Maintenance

Animals were maintained under normal growth conditions (∼20° C) on NGM agar media, with plates seeded with *E. coli* OP50 to provide nutrition. Human NRXN1, NLGN1 and SHANK3 sequences were extracted and aligned to their *C. elegans* orthologs to identify conserved variant sites as previously described^42^. The conserved missense variants were then inserted in the *C. elegans* genes via CRISPR Cas9 as previously described^42^. Briefly, Cas9 mix was reconstituted with standard materials including Cas9 nuclease, crRNA guides, tracrRNA. Short oligonucleotide templates were provided in the mix to induce mutagenesis at the desired residue. *dpy-10* crRNA was used as a co-injection marker for editing selection. Standard procedure was used for microinjections. *nrx-1, nlg-1 and shn-1* mutant strains were crossed into the *npr-1(ad609)* background for the social feeding assays. All strains used are listed in **Supplemental Table 1.**

### Food deprivation behavioral response and activity assays

Animal locomotion and activity levels were analyzed using the WorMotel device as previously described^45^. Each of the 48 wells was filled withs standard NGM agar and cooled to room temperature. For food deprivation, day 1 adult hermaphrodites were picked into a M9 bath to wash off residual bacteria and then individual animals were pipetted onto the NGM agar WorMotel wells. For fed conditions 1.5 µl of 2x concentrated OP50 was placed in each well and allowed to dry completely. Individual day 1 hermaphrodites were then picked into each well. To prevent desiccation, a 100mm petri dish containing a soaked Kimwipe was used as a humidity chamber. To assay activity, we utilized the WormWatcher automated imaging platform (Tau Scientific) to capture images at an interval of 10 seconds across an 8-hour timescale. Previously published custom MATLAB code was used to process and analyze the images^45^. In short, total pixel displacement was determined between 10 second frames and aggregated into hour bins. Image frames were manually inspected, and wells with escaped animals were removed from the final analysis. Individual replicates were combined as previously described^45^.

### Social feeding behavioral assay

To assay social feeding behavior, we utilized previously published setup of NGM-filed and OP50 seeded 6-well dishes^30,46^. To observe the behavior, 50 L4 hermaphrodites were transferred onto each OP50 seeded well with care taken to prevent bacterial buildup. A thin layer of Tween20 was added to the surface of the 6-well lid, preventing the formation of condensation during imaging. The plates are imaged using the WormWatcher automated imaging platform (Tau Scientific) for a total of 20 hours with a cluster of 10 images (over a 1-minute timecourse) being taken every hour. Quantification of social feeding aggregation was performed as previously described^30^, briefly, an animal was counted as aggregating when in contact with two or more other animals.

### Microscopy

L4 hermaphrodites were picked and aged overnight to day 1 of adulthood, and were placed on a standard 5% agarose pad, anesthetized in ∼4 μL of 100 μM sodium azide and covered with a coverslip. Animals were allowed to sit for ∼ 5 minutes to reduce subtle movement during imaging. A Leica inverted TCS SP8 laser-scanning confocal microscope equipped with a 60x objective lens was used to capture the entire head and pharynx (and underlying nerve ring). Z-stack images were taken in 0.5 microns slices, sectioning through the full Z-plane of the head. Standardization across samples was ensured using consistent laser power, zoom, and gain settings. Micrographs showing the expression pattern of NRX-1::GFP transgenes in head and nerve ring were generated in ImageJ by compressing relevant Z-slices using the standard SumStack function.

### Pathogenicity and cleavage and secretion modeling

To define the pathogenicity of the rare variants identified in humans, we utilized EMBL ProtVar and focused on two metrics defined therein. CADD (Combined Annotation Dependent Depletion) scores outline the predicted deleteriousness of single nucleotide variants^47^. We combined the CADD score with pathogenicity prediction generated by AlphaMissense^48^. To generate the CADD and AlphaMissense scores Uniprot of NLGN4X, NRXN1, and SHANK3 were obtained and the appropriate amino acid change was defined. To model the effect of NRXN1/NRX-1 variations, we utilized PrediSi to predict the probability of signal sequence cleavage and subsequent secretion^49^.

### Statistics

The number for each experiment was based on previous studies and effect size, with each experiment performed with at least 3 independent replicates and each trial performed with matched controls^30,45^. All data were analyzed and plotted in GraphPad Prism 10 and statistical significance was determined using one-way ANOVA with Tukey’s post-hoc test. Error bars on figures represent standard error of the mean (SEM) and p-values are shown in each figure to indicate significance (P<0.05).

## RESULTS

### Rare variants are conserved in *NRXN1/nrx-1*, *NLGN4X/nlg-1*, and *SHANK3/shn-1*

We identified several missense variants of interest from published studies of autism probands, including NRXN1(L18Q)^50^, NLGN4X(G>84R)^51^, NLGN4X(A283T)^51^, and SHANK3(A224T)^52^. To identify the structural location and any large-scale structural changes in the variant proteins, we utilized AlphaFold to model each protein and the impact of each variant. Surprisingly, there we no major structural modifications in the local environment, suggesting more subtle effects of these changes (**Figure 1**). To define some of these subtle impacts we modeled the variant pathogenicity using a combination of AlphaMissense^48^ and CADD^47^. Two variants, NLGN4X(A283T) and SHANK3(L143P) are predicated to be pathogenic by AlphaMissense (**Table 1**). Furthermore, all 4 variants are predicted to be likely or potentially deleterious by the CADD model, regardless of their pathogenicity status (**Table 1**).

**Table 1.** Human synaptic gene rare variants and predictions of pathogenicity.

| Human gene | Protein variant | CADD score<sup>47</sup> | AlphaMissense Score<sup>48</sup> | Variant ID |
| --- | --- | --- | --- | --- |
| NLGN4X | G84R | 23.6 (LD) | 0.2793 (benign) | ENST00000381095.8:c.250G>A |
|  | A283T | 24.7 (LD) | 0.7989 (pathogenic) | ENST00000381095.8:c.847G>A |
| NRXN1 | L18Q | 26.3 (PD) | 0.4292 (ambiguous) | ENST00000401669.7:c.53T>A |
| SHANK3 | L143P | 24.2 (LD) | 0.5747 (pathogenic) | ENST00000692848.1:c.203T>C |

\* CADD scores (LD): likely deleterious (PD) potentially deleterious
\* AlphaMissense score ranges from 0-1 (benign to pathogenic)

Alignment of the human genes with their *C. elegans* orthologs, NRX-1, NLG-1, and SHN-1 revealed strong conservation of the impacted residues (**Figure 2A, Supplemental Figure 1**). Both NLGN4X variants are located within the hydrolase functional domains, indicating potential disruption of protein activity or interactions (**Figure 2A**). Although only the two *NLGN4X* variants are situated within a functional domain of the protein (**Figure 2A**), all 4 variants carry the potential to interfere with essential protein function. NRXN1(L18Q) lies adjacent to laminin G-like domain, essential for its interaction with cognate binding partners such as Neuroligin. Similarly, the SHANK3(L143P) lies adjacent to ankyrin repeat domains, thus potentially interfering with its scaffolding function (**Figure 2A**). The location of the NRXN1(L18Q)/NRX-1(L16Q) variant lies within the cleaved signal sequence for extracellular secretion of both the human and *C. elegans* proteins. Despite the proximity to the cleavage site, the NRNX1(L18Q) variation only mildly lowered the predicted secretion of the human protein (**Table 2, Supplemental Figure 2**). In contrast, the NRX-1(L16Q) variation significantly reduced the probability of cleavage and therefore was predicted not to be secreted (**Table 2, Supplemental Figure 2**). After identifying the conserved residues for each rare variant, the variants were introduced into the respective *C. elegans* gene via CRISPR CAS9 genome editing techniques (**Figure 2B**). The variants were given allele names and are as follows – NLGN4X(G84R) is NLG-1(G59R) and *nlg-1(sy959)*, NLGN4X(A283T) is NLG-1(A261T) and *nlg-1(sy961)*, NRXN1(L18Q) is NRX-1(L16Q) and *nrx-1(sy869)*, and SHANK3(L143P) is SHN-1(L67P) and *shn-1(sy856)*.

**Figure 2.**
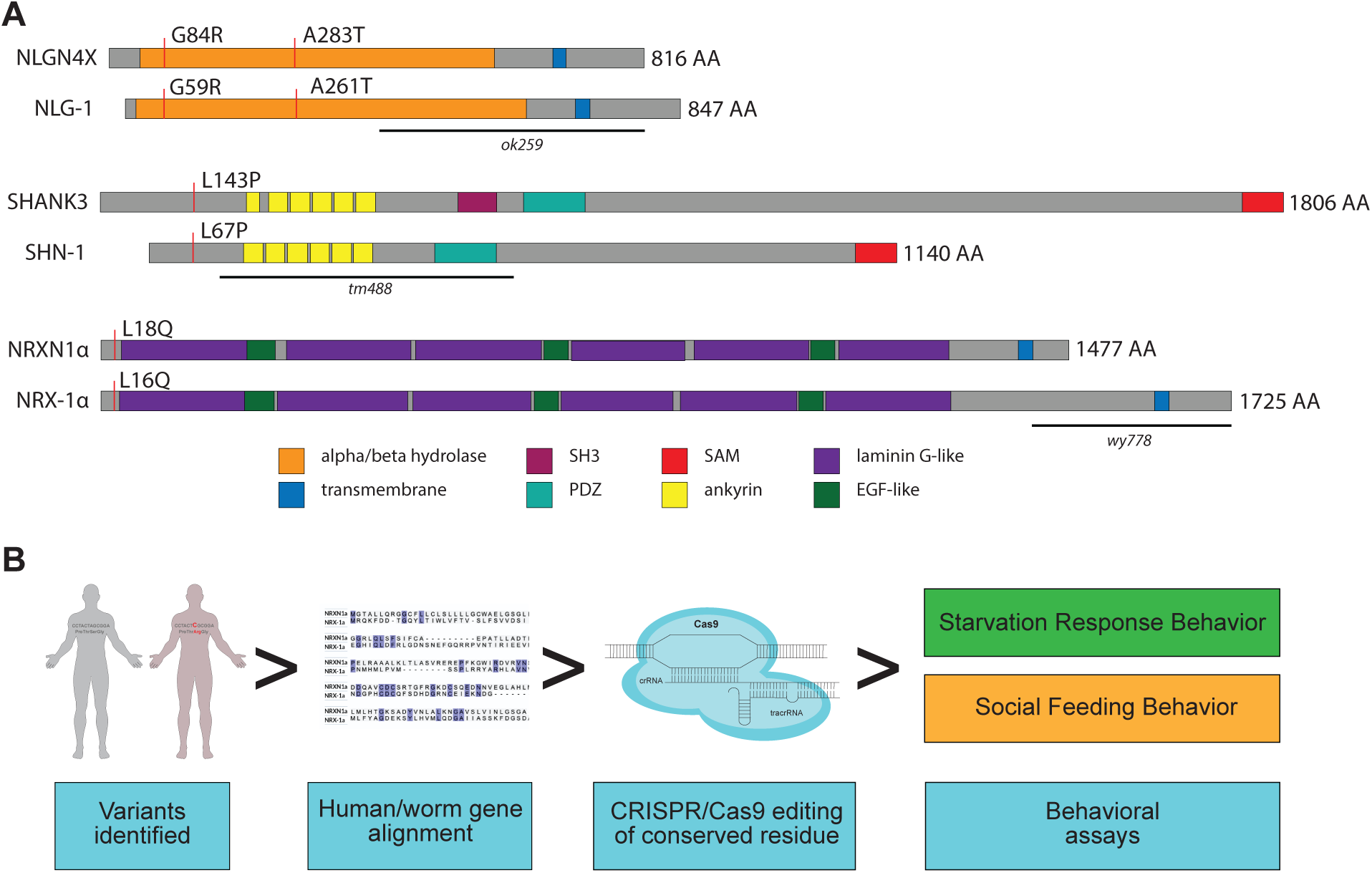
Alignment of conserved variants in NRX-1, NLG-1, and SHN-1 and workflow to test functional impact. **(A)** Human and *C. elegans* protein alignments with position of variants indicated for NRXN1(NRX-1), NLGN4X (NLG-1), and SHANK3 (SHN-1). (**B**) Cartoon of workflow of generation of variants and behavioral assays to test functional impact of controls, control loss of function alleles, and inserted conserved variants.

**Table 2.**
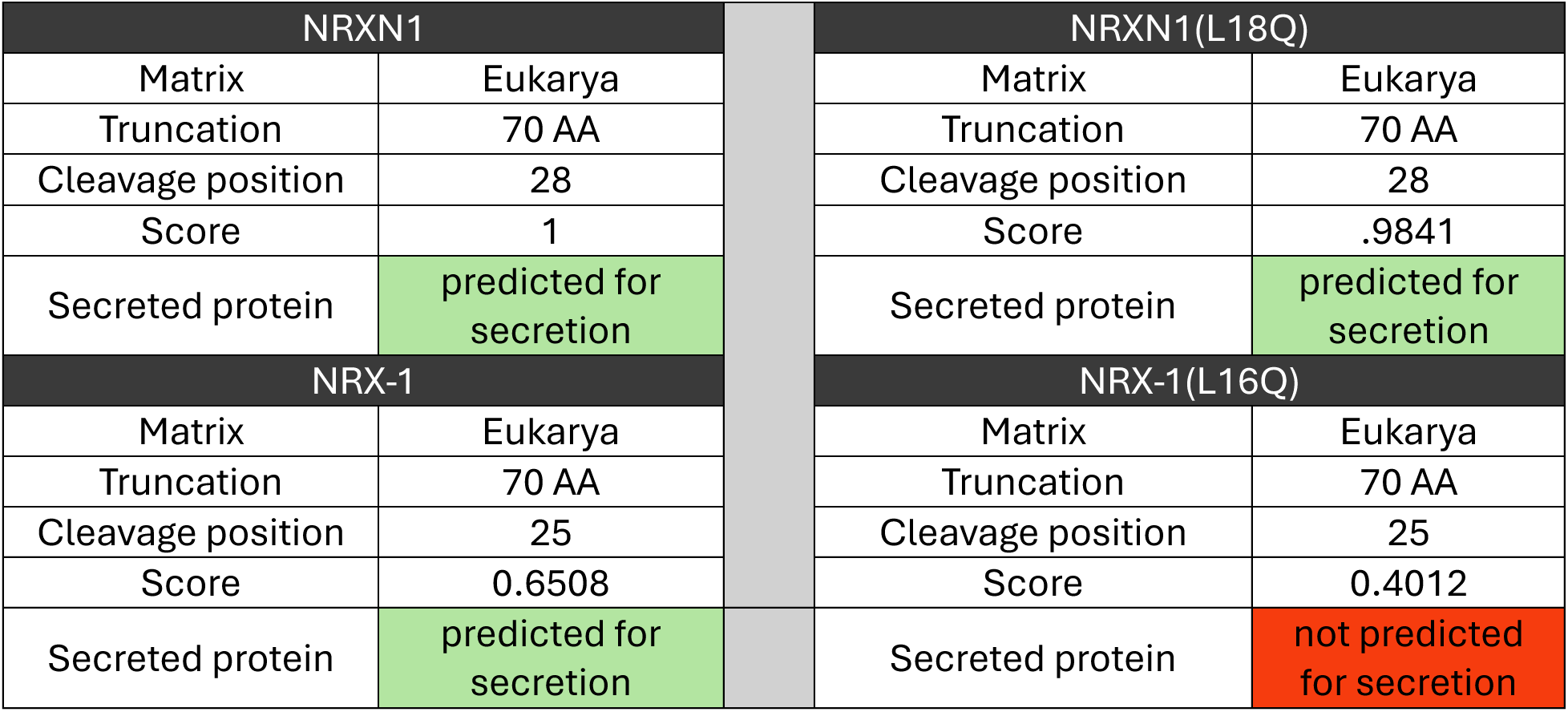
Effect of NRXN1/NRX-1 variants on protein cleavage and secretion.

| NRXN1 |  |  | NRXN1(L18Q) |  |
| --- | --- | --- | --- | --- |
| Matrix | Eukarya |  | Matrix | Eukarya |
| Truncation | 70 AA |  | Truncation | 70 AA |
| Cleavage position | 28 |  | Cleavage position | 28 |
| Score | 1 |  | Score | .9841 |
| Secreted protein | predicted for secretion |  | Secreted protein | predicted for secretion |
| NRX-1 |  |  | NRX-1(L16Q) |  |
| Matrix | Eukarya |  | Matrix | Eukarya |
| Truncation | 70 AA |  | Truncation | 70 AA |
| Cleavage position | 25 |  | Cleavage position | 25 |
| Score | 0.6508 |  | Score | 0.4012 |
| Secreted protein | predicted for secretion |  | Secreted protein | not predicted for secretion |

### Rare variants in synaptic genes alter *C. elegans* activity in response to food and food deprivation

*C. elegans* in the presence of food maintain a baseline level of activity, consistent with small movements and feeding behavior. However, when removed from food, animals significantly increase activity levels, likely to maximize foraging success. Previous work has identified several synaptic genes, including *nrx-1* and *nlr-1*, that regulate the initiation and maintenance of this hyperactivity observed with food deprivation^45,53^. Here we tested the impact of rare variants in *nlg-1*, *nrx-1*, and *shn-1* on activity levels of day 1 adult animals in the presence or absence of food. On food, *nlg-1(sy959*) and *nrx-1(sy869)* had significantly decreased activity compared to control animals (**Figure 3B**), which is in contrast to loss of function deletion alleles in *nlg-1* and *nrx-1*, which we reported to have no impact on activity levels in the presence of food^45^. We observed no impact of *nlg-1(sy961)* on activity levels of animals on food compared to controls (**Figure 3B**). Interestingly, while we observed a strong reduction of activity in the loss of function deletion *shn-1(tm488*), we saw no impact of *shn-1(sy856)* on activity levels (**Figure 3B**).

**Figure 3.**
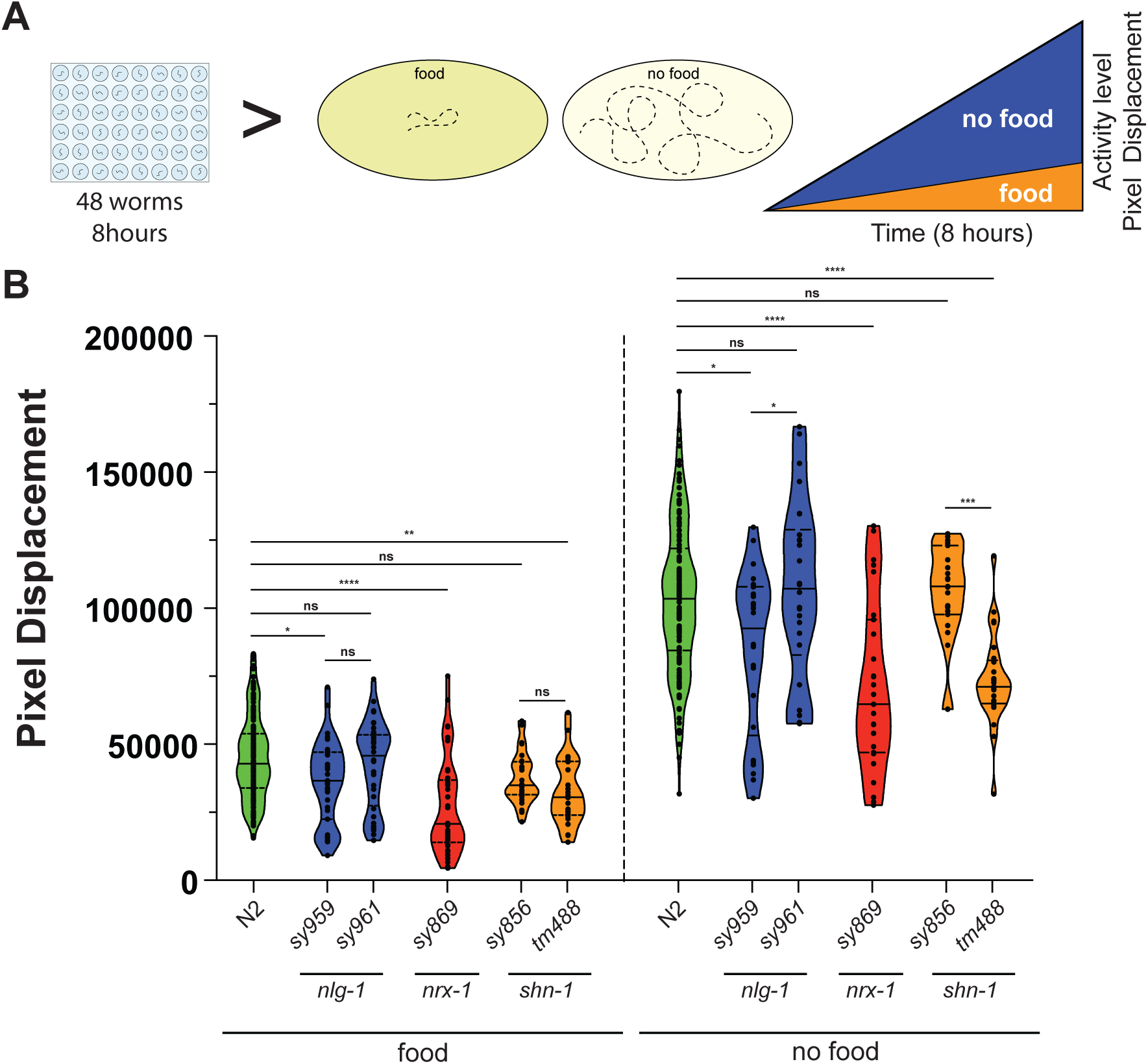
Rare variants in synaptic genes differentially alter activity levels on and off food. **(A)** Schematic of behavioral assay for activity levels of day 1 adult animals on and off food. **(B)** Average activity levels (across 8 hours) of control and synaptic gene variant animals in the presence or absence of food. *sy959* = NLG-1(G59R), *sy961* = NLG-1(A268T), *sy869* = NRX-1(L16Q), *sy856* = SHN-1(L67P), *tm488* = large deletion loss of function *shn-1*. Dots represent individual animals. Statistical significance was determined by One-way ANOVA with Tukey posthoc test.

In response to food deprivation, we observed the expected increase in activity levels of control N2 animals (**Figure 3A-B, Supplemental Figure 3 A-E**). *nlg-1(sy959)* animals off food showed a reduction in activity compared to controls, again in contrast to previous findings for loss of *nlg-1*^45^, while *nlg-1(sy961)* had no impact on activity levels off food compared to controls (**Figure 3B**). Similar to loss of *nrx-1*, *nrx-1(sy869)* animals had reduced activity levels when food deprived compared to controls (**Figure 3B**). Consistent with activity levels on food, loss of *shn-1* also showed a reduction in activity levels compared to controls, while *shn-1(sy856)* had no impact on activity levels off food (**Figure 3B**). Therefore, we find a surprising difference comparing the impact of rare variants to loss of function deletion alleles in both *nlg-1* and *shn-1* alleles on behavioral responses to food and loss of food (**Figure 3B**, **Table 3**). These results indicate that some of the rare variants likely involve loss of function, but with some also having novel functional impacts compared to loss of the genes, indicating potential gain of function or dominant negative impact.

**Table 3.** Summary and comparison of behavioral changes observed with synaptic gene variants.

| Gene variant | Change | Activity |  | Feeding |  |
| --- | --- | --- | --- | --- | --- |
|  |  | on food | off food | solitary | social |
| *nlg-1(ok259)* | deletion | no effect | no effect | no effect | decrease |
| *nlg-1(sy959)* | G59R | decrease | decrease | no effect | no effect |
| *nlg-1(sy961)* | A268T | no effect | no effect | no effect | decrease |
| *nrx-1(wy778)* | deletion | no effect | decrease | no effect | decrease |
| *nrx-1(sy869)* | L16Q | decrease | decrease | increase | decrease |
| *shn-1(tm488)* | deletion | decrease | decrease | no effect | decrease |
| *shn-1 (sy858)* | L67P | no effect | no effect | no effect | decrease |

\* lighter color blocks indicate result come from previous studies

### Rare variants in synaptic genes differentially alter solitary and social feeding behaviors

We next tested the impact of the rare variants on social feeding behaviors of *C. elegans*. Wild isolate strains of *C. elegans* and other related *Caenorhabditis* species can feed alone or in groups, with many strains showing a range of either solitary or social feeding behavior, where animals group together in aggregates in the presence of food^54,55^. The *C. elegans* control lab strain (N2 Bristol) displays solitary feeding, where animals avoid one another when on a food lawn or bacterial substrate ^55^. Point mutation or loss of function alleles in the *npr-1* gene (orthologue of neuropeptide Y receptor), induce social aggregation feeding behavior^55^. Here, we quantified aggregating animals to quantify social feeding behaviors in animals with rare variants in *nlg-1*, *nrx-1*, and *shn-1* alone (solitary background) or with *npr-1(ad609)* allele (social background).

Compared to the control N2 solitary feeding animals, which show almost no aggregation, we observed no impact of the rare variants in *nlg-1*, *shn-1*, or of the *shn-1* loss of function deletion allele (*tm488*) (**Figure 4B-C**). However, *nrx-1(sy869)* significantly increased aggregation in the solitary background (**Figure 4B-C**). Interestingly, this impact on behavior is unique to the *nrx-1(sy869*) variant, as there is virtually no aggregation observed in other *nrx-1* alleles (eg. *nrx-1(wy778*))^30^ in solitary genetic backgrounds. The same variants in the social *npr-1(ad609)* background showed different impacts, where we observed a strong reduction in aggregation with *nlg-1(sy961)*, and *shn-1(sy856*) (**Figure 4B-C**). This is comparable to the reduction observed in social feeding behavior of *nrx-1* and *nlg-1* loss of function deletions (**Figure 4B-C**)^30^, while *nlg-1(sy959)* had no impact on aggregation. We observed a reduction in aggregation with loss of *shn-1(tm488)*, indicating a role for *shn-1* in social feeding behavior in *C. elegans* (**Figure 4B-C**). Interestingly, the rare *shn-1* variant had a stronger impact on aggregation and social feeding than the loss of function deletion (**Figure 4B-C**). Overall, the reductions in aggregation in the social background with some rare variants in *nrx-1* and *nlg-1* are similar to previously observed loss of these genes, which fine tune neuronal connections and the activity within the RMG circuit that drives social feeding behaviors^30^. Remarkably, the rare variant in *nrx-1, sy869,* induces some social feeding in a solitary background, further indicating the potential for *nrx-1* to tune solitary and social feeding behaviors.

**Figure 4.**
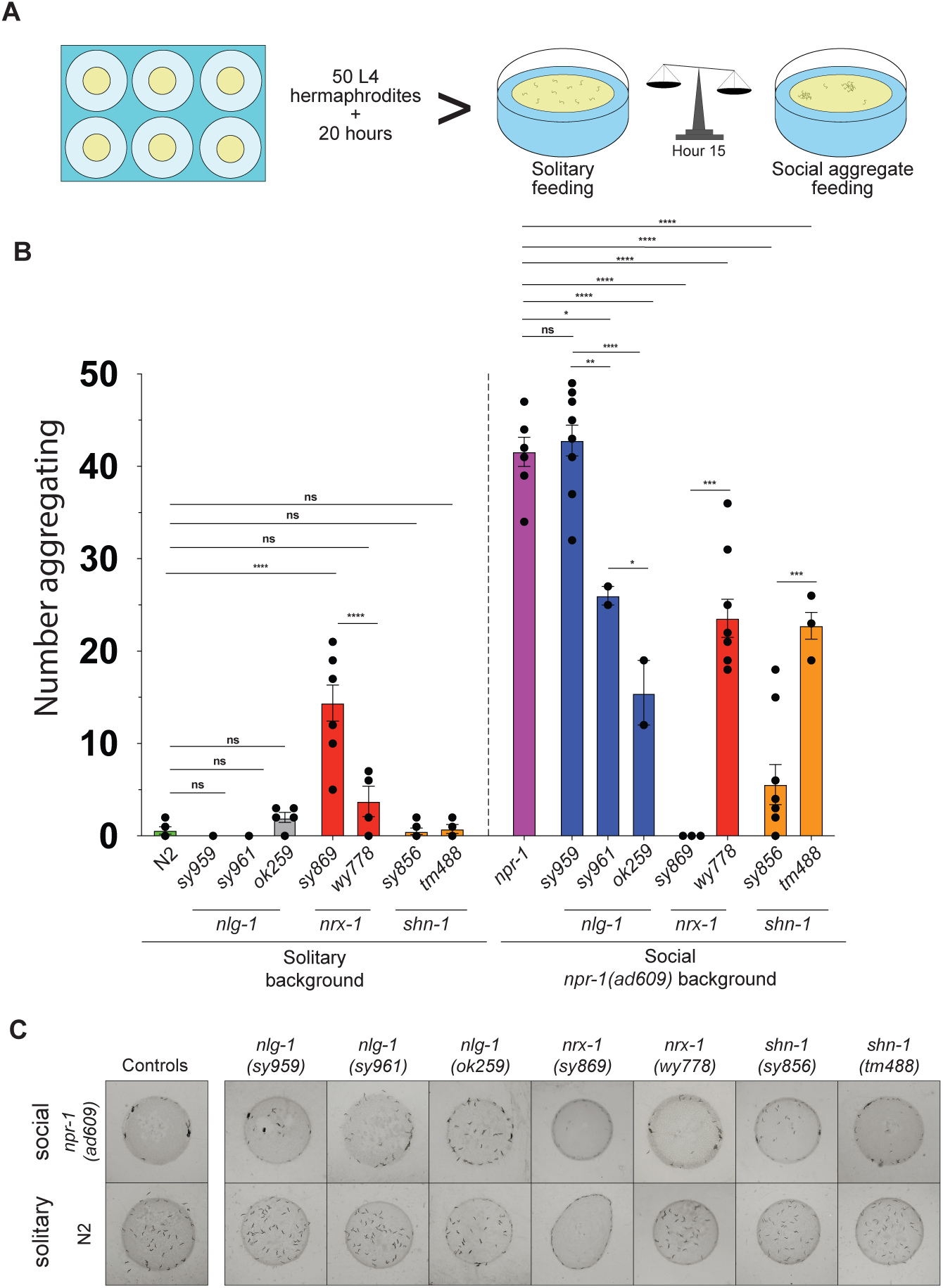
Rare variants in synaptic genes impact solitary and social feeding behaviors. **(A)** Cartoon of solitary and social feeding behaviors and assay. **(B)** Number of day 1 adult aggregating animals in the solitary (left) or social (*npr-1(ad609)*) (right) background out of 50 animals total. Dots represent replicates. Statistical significance was determined by One-way ANOVA with Tukey posthoc test. **(C)** Representative images of each variant and controls in solitary and social (*npr-1(ad609)*) genetic backgrounds.

### NRX-1 L16Q does not impact protein expression or localization, but alters behavior

Given the novel behavioral phenotypes we observed with the rare *nrx-1(sy869)* variant, we sought to further characterize the impact of the variant on NRX-1. To do this, we created expression plasmids with the long isoform of NRX-1 cDNA tagged with superfolder GFP^45^. We made versions with or without the NRX-1(L16Q) variant driven in all neurons using the *ric-19* promoter (*ric-19p::sfGFP::nrx-1* and *ric-19p::sfGFP::nrx-1(L16Q)*). The sfGFP tag was inserted downstream of the predicted signal sequence and cleavage site, including the rare variant, and upstream of the first LNS domain. The expression plasmids were then used to create transgenic animals with extrachromosomal arrays in a *nrx-1(wy778)* loss of function deletion background. Consistent with previous studies expressing *C. elegans* NRX-1 and human NRXN1^45,56^, we observed GFP expression in the nerve ring of day 1 adult animals, which was comparable in expression level and gross localization regardless of the L16Q variant (**Figure 5A**). This suggests that although the variant is in the predicted signal sequence and could interfere with cleavage of the signal sequence, the protein is still expressed and localized. We next tested the impact of the transgenes on the food deprivation response behavior to analyze functional impact of the variant in the transgene. Without food, expression of wildtype NRX-1 in all neurons increased activity levels compared to loss of *nrx-1(wy778)* (**Figure 5B**), suggesting partial restoration of *nrx-1* function and the behavioral response to food deprivation, as previously reported^45^. Interestingly, expression of NRX-1(L16Q) in all neurons, significantly decreased activity levels off food compared to controls, and of the *nrx-1(wy778)* loss of function deletion (**Figure 5B**). Therefore, although the L16Q variant does not appear to reduce expression or change localization of the NRX-1 protein, it is unable to restore NRX-1 function or the behavioral response to food deprivation, and may have some gain of function, further disturbing the behavioral phenotype. Lastly, we wanted to test the impact of the transgenes on solitary and social feeding behaviors. In contrast with insertion of the *nrx-1(sy869)* variant in the genome, neither the NRX-1 or NRX-1(L16Q) transgene significantly increased aggregation levels compared to controls (**Figure 4C**).

**Figure 5.**
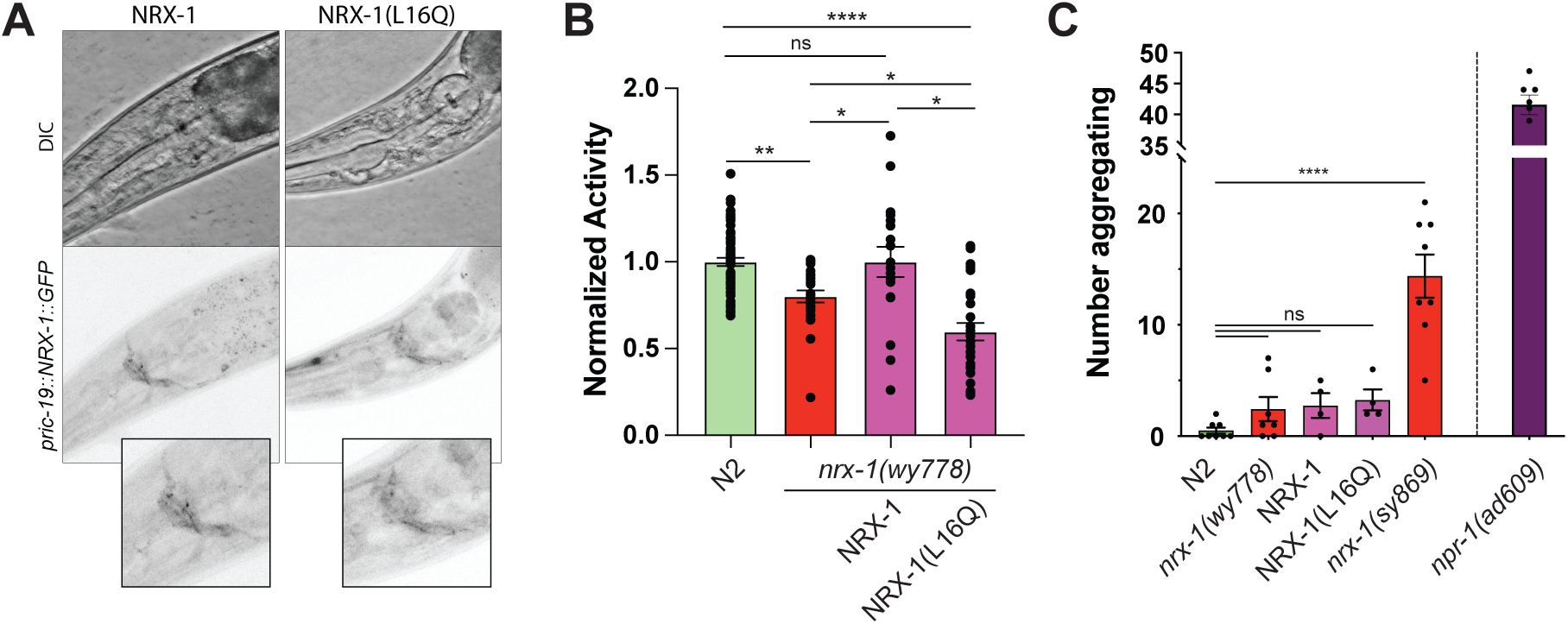
Expression of NRX-1(L16Q) disrupts the behavioral response to food deprivation. **(A)** Representative images showing expression and localization of NRX-1 and NRX-1(L16Q) in all neurons (driven with *ric-19* promoter) tagged with superfolder GFP. Insets highlight expression in the nerve ring. **(B)** Activity levels of day 1 adults without food of controls, and *nrx-1(wy778)* animals alone, with NRX-1, or NRX-1(L16Q). Dots represent individual animals. Statistical significance was determined by One-way ANOVA with Tukey posthoc test. **(C)** Aggregation of NRX-1 and NRX-1(L16Q) transgenes in solitary background. Data from N2, *nrx-1(wy778*), *nrx-1(sy869)* and *npr-1(ad609)* was replotted from Figure 3 to compare to the transgenes.

## DISCUSSION

Uncovering the genetic mechanisms driving the diverse behavioral changes in autism and related conditions is especially difficult due to the combinatorial effects of the many genes involved, multigenic risk factors, and environmental involvement. In this work, we bypassed some of these challenges by studying genetic drivers of autism in a tractable model, harnessing the simple neuronal biology, circuitry, and behaviors of *C. elegans* to analyze clinically identified variants in *NRXN1*, *NLGN4X*, and *SHANK3*. In contrast with previous characterization of large deletion and loss of function alleles^39,57–64^, we focused on understanding the impact (and potential pathogenicity) of rare missense variants leading to single amino acid substitutions in well conserved residues of the human and *C. elegans* genes. Overall, we observed significant behavioral phenotypes in NRX-1, NLG-1, and SHN-1 across two well-characterized behavioral paradigms, social feeding behavior and response to food/food deprivation (**Table 3**). Although many of the variants displayed phenotypes consistent with loss or partial loss of function, we also observed several interesting differences comparing rare variants and respective loss of function alleles. As an example, in contrast to the *nlg-1*(*ok259*) putative null allele, we see a significant impact of the two *nlg-1* variant alleles depending on the behavior, with *nlg-1(959)* having a strong loss of function phenotype only in the food/food deprivation assay and *nlg-1(S61)* having a strong loss of function only in social feeding behavior. In addition, we observed novel and strong loss of function phenotypes across both food/food deprivation response and social feeding behavior in the *shn-1(tm3488)* putative null allele. Further highlighting phenotypic variation between alleles, *shn-1(sy856)* only significantly impacted social feeding behavior suggesting a more nuanced impact of the missense variation on protein function within this neuronal circuit. Lastly, and of particular interest, *nrx-1(sy869)* had a significant (and opposite) pattern across the two behavioral assays, displaying a loss of function phenotype in response to food deprivation (consistent with LOF alleles), but a gain of function phenotype in solitary feeding behavior (**Table 2**). Furthermore, we observe evidence of a complex epistatic interaction between *npr-1* and *nrx-1(L16Q)* in that the combination seems to eliminate the aggregation inherent to both single mutants. This context dependent switch in behavioral phenotypes suggests a subtle impact of the conserved residue in the function of the protein. Furthermore, this suggests a potential difference in the functional role or perhaps a change in physical interactions for NRX-1 among the two behavioral circuits. Taken together with the robust phenotypes we observed in several of the variants, our work defines the contribution of these variants to the mechanism of protein dysfunction.

While individual variants in synaptic genes, including both large deletions and single nucleotide changes, have long been associated with neurodevelopmental conditions, we still lack a tractable genetic model in which to test the effects across many genes, variants, and behaviors. Genes like those in the neurexin family pose a particularly cumbersome challenge in mammalian and cell culture models given the nature of a multi-gene family and the differential generation of thousands of isoforms, some of which likely play highly specialized and context-dependent roles. When analyzing the differences in functional impacts between gene deletions (loss of function or null) and single amino acid substitutions there are many things to consider. In the most straightforward case, a large deletion eliminates the production of the wildtype protein, and if a truncated mutant protein is produced the levels are likely diminished compared to wildtype levels. Introducing complexity to the genetic locus through isoform production can increase the chance that a truncated (but still functional) product is produced even the event of a large deletion.

Further complicating matters, complex genetic loci such as those that generate the neurexins undergo significant regulation at the level of splicing meaning that disruptions in splicing sites of junctions can introduce rare exon variations leading to protein dysfucntion^65,66^. In comparison, missense variants have more unpredictable effects, ranging from benign to evoking structural or domain changes and causing pathogenicity. Despite the potential for deleterious impact on protein and neuronal function, it remains difficult to accurately predict the full effect of single residue variations. Our findings in comparing multiple genes and variants across multiple behaviors, highlight the distinct impact of variants across different circuits and behaviors. A single variant can induce loss of function in one behavior, but have no impact on another behavior. For genes like neuroligins and neurexins, which have many isoforms that can have neuron- and synaptic-specific functions, a variant can not surprisingly impact only a subset of isoforms, functions, and behaviors. Our results demonstrate the need to analyze multiple readouts of gene function, in this case behaviors, when defining the impact of variants in synaptic genes. Studying only a single behavioral assay would miss the functional impact of some gene and variants. Further, the comparison across behaviors and genes, in comparison with known loss of function alleles and variants, allows mechanistic insight into gene function in the generation of behaviors.

Recent advances in structural and pathogenicity modeling of proteins and protein variants allow predictions in whether individual missense variants will impact the function of the protein based on conservation and structural changes. However, predicted pathogenicity does not always correlate with actual function when tested or with impact in behavior. For example, modeling of the NLGN4X(G84R) conserved residue revealed a predicted benign impact via the pathogenicity model, which stands in stark contrast with the variants’ significant impact in food/food deprivation response behaviors. Likewise, NRXN1(L18Q) is predicted to have a functionally ambiguous impact, while the variant clearly has impact on gene function and multiple behaviors. Further, the NRX-1(L16Q) variant was predicted to change the signal sequence and cleavage, decreasing secretion to the membrane, which we did not observe when we expressed the variant protein. Perhaps the variant alters cleavage, but not secretion, and the signal sequence somehow alters a function or interaction of the protein. The NRXN1(L18Q) variant was predicted to have a milder impact than the C. elegans variant, but perhaps the variant lowers efficiency of the cleavage, and may result in retention of the signal sequence, with similar impact. Overall, although the existing predictive models can inform on potential for pathogenicity, there is still a significant need to couple computational modeling and functional assays of gene function (genetic/molecular and behavioral assays) to define the functional impact of a missense mutation. Together, our work contributes to the growing number of studies that define the impact of genes and rare variants associated with neuropsychiatric and neurodevelopmental conditions through study in genetically tractable models with small neuronal circuits. Here, we provide evidence that *C. elegans* represents an increasingly powerful tool to link specific protein variations across multiple genes to changes in neuronal function and ultimately alterations in multiple behaviors.

## Data Availability Statement

All data and materials are available upon request to the corresponding author. All data are available in the main text or the supplementary materials and source data are provided in the Source Data file.

## Acknowledgements

The authors thank Paul Sternberg and Sandy Wong for creating and sharing the rare variant strains. We also thank members of the Hart lab for comments on the manuscript. Some strains were provided by the CGC, which is funded by NIH Office of Research Infrastructure Programs (P40 OD010440). This work was supported in part by Penn ASPE, SFARI BTI award (MPH), NIH R35GM146782 (MPH), and NIH R56MH096881(MPH).

## Author Contributions

DH, BLB, and MPH conceived and designed the study and experiments, and DH, WRH, MG, and BLB conducted all experiments. DH processed, analyzed, and interpreted all data, with help from MG and BLB. DH wrote the manuscript with assistance from MPH, and all authors reviewed, revised, and approved the manuscript.

---

**Supplemental Table 1.** *C. elegans* strain information.

| Gene | Strain | Allele | Source |
| --- | --- | --- | --- |
| *nrx-1* | TV13570 | *nrx-1(wy778)* | CGC |
|  | MPH49 | *nrx-1(wy778);npr-1(ad609)* | previous work<sup>30</sup> |
|  | PS7330 | *nrx-1(sy869)* | this study |
|  | MPH219 | *nrx-1(sy869);npr-1(ad609)* | this study |
|  | MPH224 | *nrx-1(wy778); hpmEx36[ric-19p::sfgfp::nrx-1($\alpha$); lin-44::gfp]* | this study |
|  | MPH225 | *nrx-1(wy778); hpmEx37[ric-19p::sfgfp::nrx-1($\alpha$)(L16Q); lin-44::gfp]* | this study |
|  | MPH226 | *hpmEx36[ric-19p::sfgfp::nrx-1($\alpha$); lin-44::gfp]* | this study |
|  | MPH227 | *hpmEx37[ric-19p::sfgfp::nrx-1($\alpha$)(L16Q); lin-44::gfp]* | this study |
| *nlg-1* | PS7711 | *nlg-1(sy959)* | this study |
|  | MPH220 | *nlg-1(sy959);npr-1(ad609)* | this study |
|  | PS7713 | *nlg-1(sy961)* | this study |
|  | MPH221 | *nlg-1(sy961);npr-1(ad609)* | this study |
|  | VC228 | *nlg-1(ok259)* | CGC |
|  | MPH43 | *nlg-1(ok259);npr-1(ad609)* | previous work<sup>30</sup> |
| *shn-1* | KP7032 | *shn-1(tm488)* | previous work<sup>67</sup> |
|  | MPH222 | *shn-1(tm488);npr-1(ad609)* | this study |
|  | PS7259 | *shn-1(sy856)* | this study |
|  | MPH223 | *shn-1(sy856);npr-1(ad609)* | this study |

**Supplemental Figure 1.**
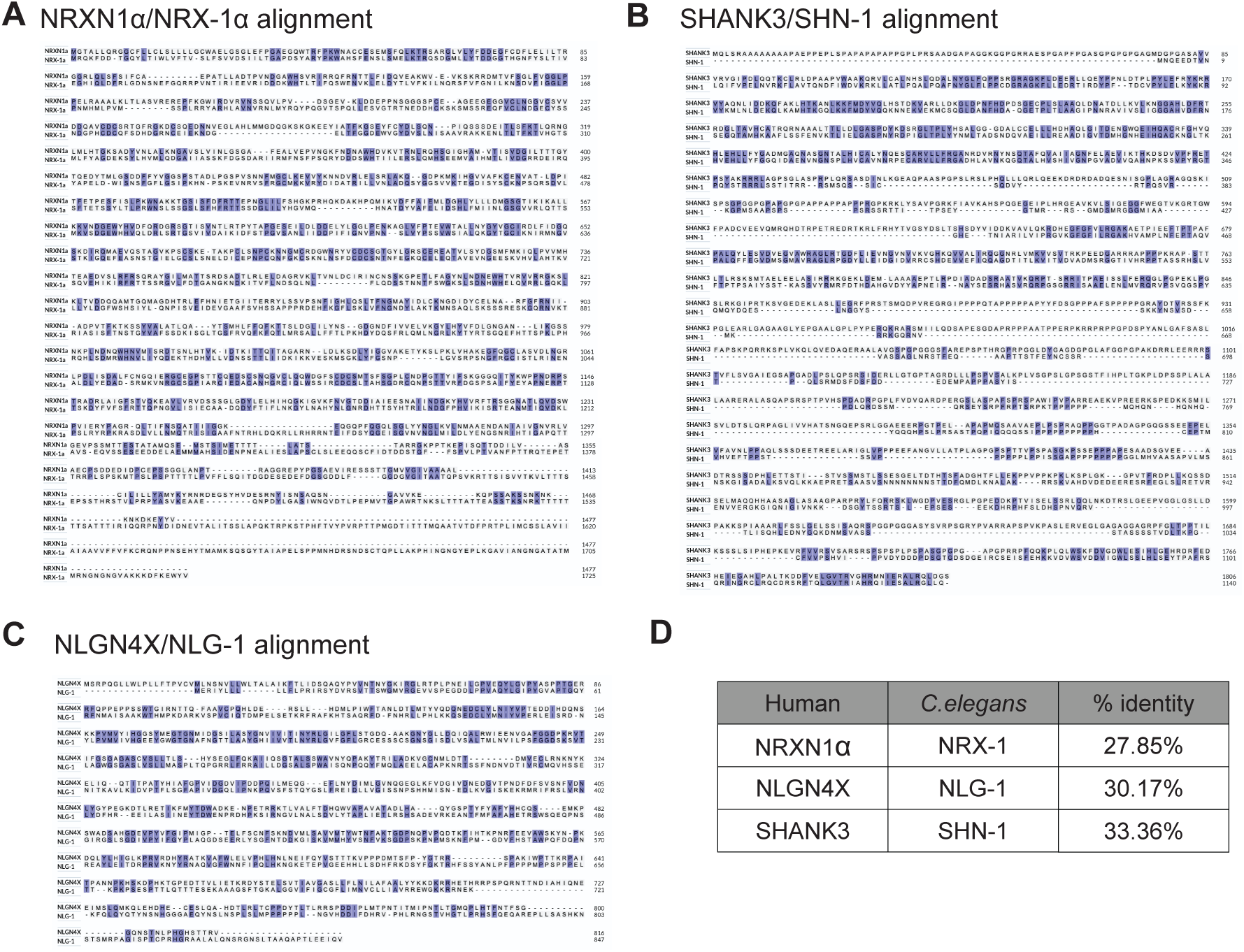
Clustal Protein alignments of synaptic genes. Longest isoform sequences were obtained for NRXN1, NlGN4X, and SHANK3 along with their C elegans orthologues. Protein alignments were done utilizing EMBL Clustal Omega Multiple Sequence Alignment tool.

**Supplemental Figure 2.**
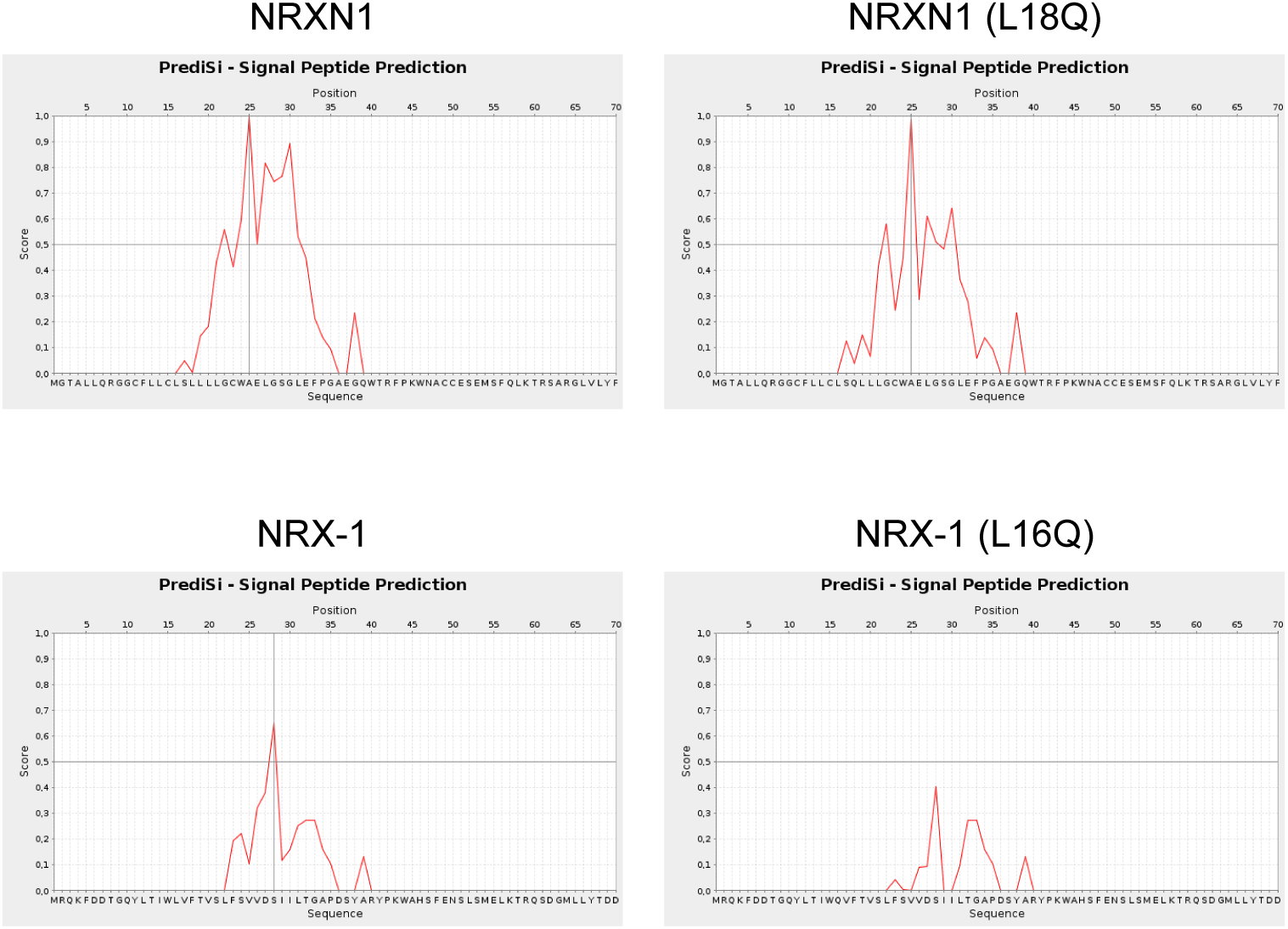
PrediSi predictions for NRXN1 and NRX-1 proteins and impact of rare variant. 70 AA truncated proteins were analyses for the probability and location of cleavage and prediction of extracellular secretion.

**Supplemental Figure 3.**
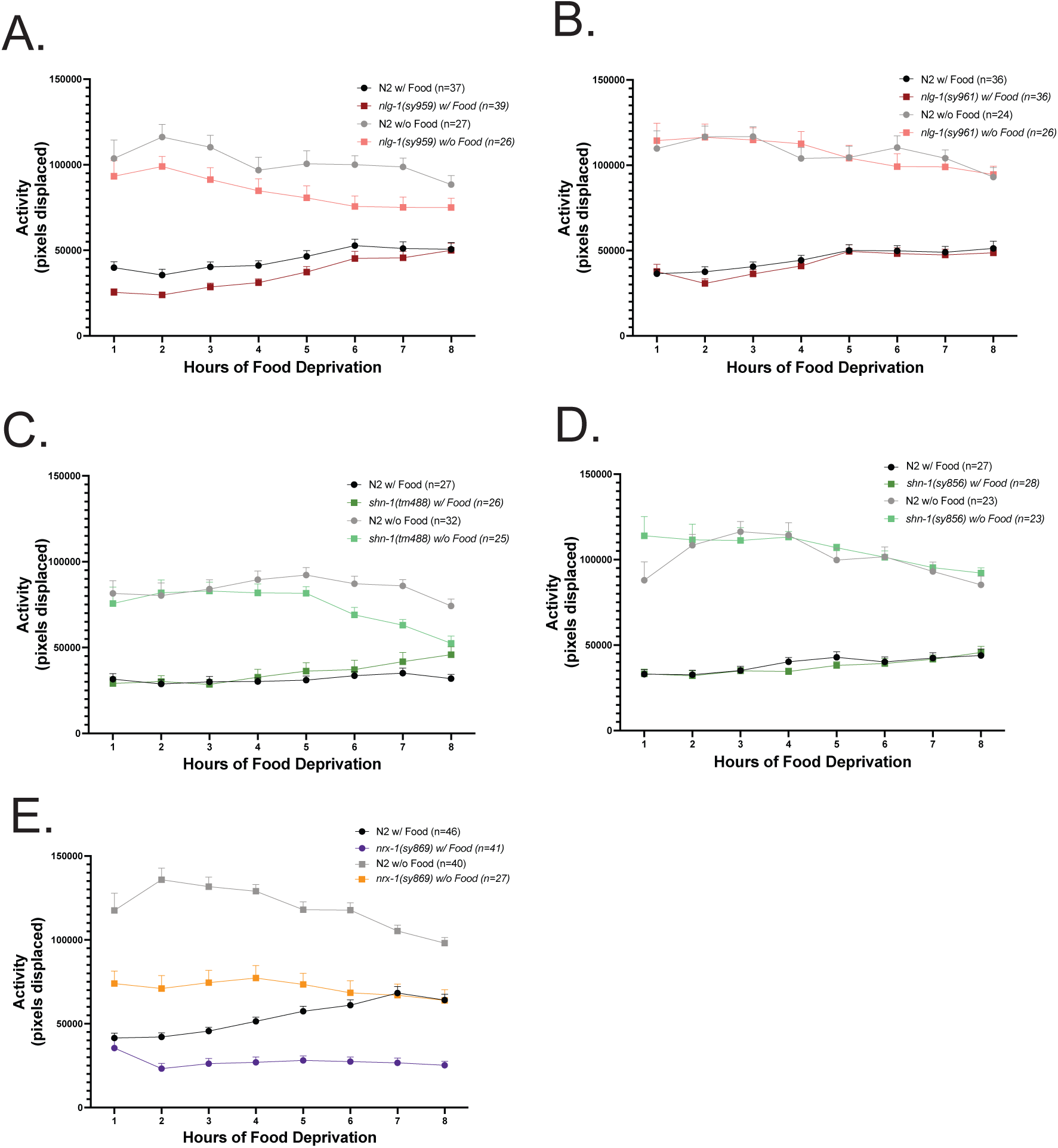
Activity (over eight hours) of controls and synaptic gene variants. Average activity levels (across 8 hours) of control and synaptic gene variant animals in the presence of food and upon food deprivation. **(A)** *nlg-1(sy959)* **(B)** *nlg-1(sy961)* **(C)** *shn-1(tm488)* **(D)** *shn-1(sy856)* **(E)** *nrx-1(sy869)*. Dots represent the average activity value across individuals. Statistical significance was determined by One-way ANOVA with Tukey test correction.

